# Cryopreservation of the collector urchin embryo, *Tripneustes gratilla*

**DOI:** 10.1101/2024.01.22.576601

**Authors:** Charley E. Westbrook, Jonathan Daly, Brian Bowen, Mary Hagedorn

**Author notes:** Corresponding author. Charley Westbrook Hawaii Institute of Marine Biology, University of Hawaii at Manoa, 46-007 Lilipuna Rd., Kaneohe, HI, 96744, USA *E-mail Address:.

## Abstract

The collector urchin, *Tripneustes gratilla*, is an ecologically important member of the grazing community of Hawaii’s coral reefs. Beyond its ability to maintain balance between native seaweeds and corals, *T. gratilla* has also been used as a food source and a biocontrol agent against alien invasive algae species. Due to overexploitation, habitat degradation, and other stressors, their populations face local extirpation. However, artificial reproductive techniques, such as cryopreservation, could provide more consistent seedstock throughout the year to supplement aquaculture efforts. Although the sperm and larvae of temperate urchins have been successfully cryopreserved, tropical urchins living on coral reefs have not. Here, we investigated the urchin embryos’ tolerance to various cryoprotectants and cooling rates to develop a cryopreservation protocol for *T. gratilla*. We found that using 1M Me2SO with a cooling rate of 9.7°C/min on gastrula stage embryos produced the best results with survival rates of up to 85.5% and up to 50.8% maturation to the 4-arm echinopluteus stage, assessed three days after thawing. Continued research could see cryopreservation added to the repertoire of artificial reproductive techniques for *T. gratilla*, thereby assisting in the preservation of this ecologically important urchin, all while augmenting aquaculture efforts that contribute to coral reef restoration.

## Introduction

Herbivorous guilds exert a particularly strong influence on the benthic species assemblage of reefs [47,73]. Excessive algae growth can smother reefs by outcompeting benthic species for space and resources. An abundant and diverse grazing community can efficiently limit algae blooms, in turn promoting coral growth and propagation [12,34]. Discussions of herbivorous guilds on coral reefs tend to highlight the importance of grazing fish communities [47,69,9], but often omit the complicit role that macroinvertebrates play in removing fleshy algae [41,24]. Urchins, for example, provide functional redundancy to the grazing pressure of herbivorous fish, in some cases even superseding fish browsing on overfished, depauperate reefs [17,45]. For many reefs, sea urchins function as keystone species by providing indispensable grazing pressure [34,16,17,10,60], contributing much of the resiliency that safeguards coral-dominated benthic communities from phase shifts to algal-dominated communities [6,45,79]. Across broadly distributed reefs, the tropical sea urchin, *Tripneustes gratilla*, is an essential primary grazer involved in the recycling of nutrients and biodiversity maintenance [42,43]. The collector urchin, *T. gratilla*, has a generalist grazing behavior that enables it to effectively reduce algal cover from reefs, fulfilling the role of ecosystem engineer by freeing up space from macrophytes and allowing other species to settle on the reefs [68,73].

Unfortunately, *T. gratilla* populations are afflicted by a variety of anthropogenic pressures including over exploitation, diminishing habitat size, the mooring and anchoring of motorboats along with various other stressors related to coastal development [23,38]. Urchin populations from around the world have also manifested vulnerabilities to waterborne pathogens, to which, several populations have succumbed without recovery [46,19,22,79,78].

Although cryopreservation of reproductive material has been achieved for members of the Echinoidea, such as *Strongylocentrotus intermedius* [4], *Tetrapigus niger* and *Loxechinus albus* [7], *Evechinus cloroticus* [2], *Paracentrotus lividus* [56], *Echinometra Lucunter* [15], investigating artificial reproductive techniques for *T. gratilla* has yet to include such novel techniques as cryopreservation for their sperm, eggs or their embryos. Cryopreservation is a growing field of research with potential applications in conservation, and aquaculture [54,55]. The cryopreservation of sea urchin gametes and larvae could provide high quality reproductive material all year round, thus allowing the aquaculture of these organisms to be independent of seasonality or natural reproductive cycles [26,1,56]. Additionally, applications of cryopreserved material could be expanded beyond supplementing the production of seedstock in the aquaculture industry, to include hybridization, selective breeding, increasing the availability of certain lines, developing and maintaining genetically modified stocks, a tool for ecotoxicological studies, as well as providing propagules for assisted gene flow [26,1,56,8, 55, 31]. Cryopreservation can also be applied to the protection of genetic diversity and the conservation of endangered species [72,33].

Marine invertebrate embryos and larvae can be the most fragile stage in the animals’ life history, making them particularly vulnerable to pollutants, osmotic and thermal stress [21,76,37,50,59,8,57], this in turn increases the challenge of cryopreserving them successfully. In order to elucidate potential cryopreservation protocols that could provide reliable means of banking and preserving *T. gratilla* larvae, several experiments were performed. A toxicity analysis was conducted with a variety of cryoprotectant agents, at different concentrations, to gauge the collector urchin larvae’s initial sensitivity to cryoprotectant agent exposure. Based on candidate cryoprotectants screened from these toxicity experiments, cryopreservation protocols were tested at various cooling rates. The most effective cryopreservation protocol was then implemented at a variety of *T. gratilla*’s ontogenetic stages, to determine if any of its early developmental phases were more or less receptive to the protocol.

## Methods

### Animal Maintenance and Reproduction

Adult sea urchins (*T. gratilla*) were collected along the southern (Sand Island) and western (Kahe Point) shores of O’ahu, Hawai’i, under Special Activities Permits Number SAP 2018-03, SAP 2019-16, SAP 2020-25, and SAP 2020-2021. Animals were transported to the laboratory wrapped in wet towels in a cooler, and kept in flow through tanks with aerated seawater at ambient temperature (25-27°C). Sea urchins were fed the red algae *Gracilaria salicornia*.

Prior to experimentation, the study animals were fasted for three days in order to empty waste from their guts prior to gamete collection. Spawning was induced by gently shaking the urchins until gametes were released. This process did not harm the urchins. Individual females (see Table 1 for details) were placed upside down on a beaker, and their eggs were released into a volume of 100 ml of 0.2 µm-filtered seawater (FSW) (Merck Millipore, Billereca, MA, USA) and held at room temperature (23–25°C). Male urchins (see Table 1 for details) were gently blotted dry to remove any excess seawater and then their sperm was collected neat (or dry) by drawing the gametes directly from their gonopores with a pipette and transferring the samples to individual Eppendorf tubes on ice. The goal was to keep the sperm in as concentrated a condition (∼10^10^ cells/ml upon spawning) as possible as it increased the longevity of the sperm.

**Table 1:** Summary of the number of treatments per experiment, number of females and number of pooled males’sperm used for fertilization to create the larvae used in the treatment and the number of eggs per treatment

| *Experiment* | *# Treatments* | *# Females* | *# Pooled Males* | *# Larvae/*<br>*Treatment* |
| --- | --- | --- | --- | --- |
| *Toxicity* | 10 | 6 | 3 | ~100 |
| *Permeant CPAs*<br>*and Cooling Rates* | 12 | 8 | 3 | ~100 |
| *Trehalose* | 4 | 8 | 3 | ~100 |
| *Cryo Protocol* | 5 | 8 | 3 | ~100 |
| *Totals* | 31 | 30 | 12 | 22,800 |

Egg quality was assessed by visual microscopic inspection to ensure they were uniformly round and undamaged prior to experimentation. Sperm motility was assessed for each male using computer assisted sperm analysis [80], to only include sperm with progressive motility rates greater than 70%. Eggs from each female were artificially fertilized with sperm pooled from three different males. Fertilization was conducted with a sperm to egg ratio of ≤ 2500:1 for these experiments, as recommended by the EPA’s fertilization assessment for *T. gratilla* [74]. Eggs were held for an hour to fertilize before being rinsed with FSW, after which time, they were transferred to a 1-L Pyrex bowl filled with FSW and incubated at 27°C.

Figure 1 identifies the order of experiments for development of urchin embryo cryopreservation. Toxicity trials exposed the embryos to a variety of permeating and non-permeating cryoprotectant agents (CPAs) at different concentrations at one single developmental time point (gastrula stage) to determine their effect over time. The least toxic cryoprotectant agents from these trials were then used in the following cryopreservation trials to determine optimal freezing rates for the urchin larvae. The non-permeant cryoprotectant agent trehalose was then added to most effective protocol from the previous experiment, to assess if it improved survival rates. Lastly, the optimal cryopreservation protocol was applied to various developmental stages to determine maximal post-thaw survival and development.

**Figure 1:**
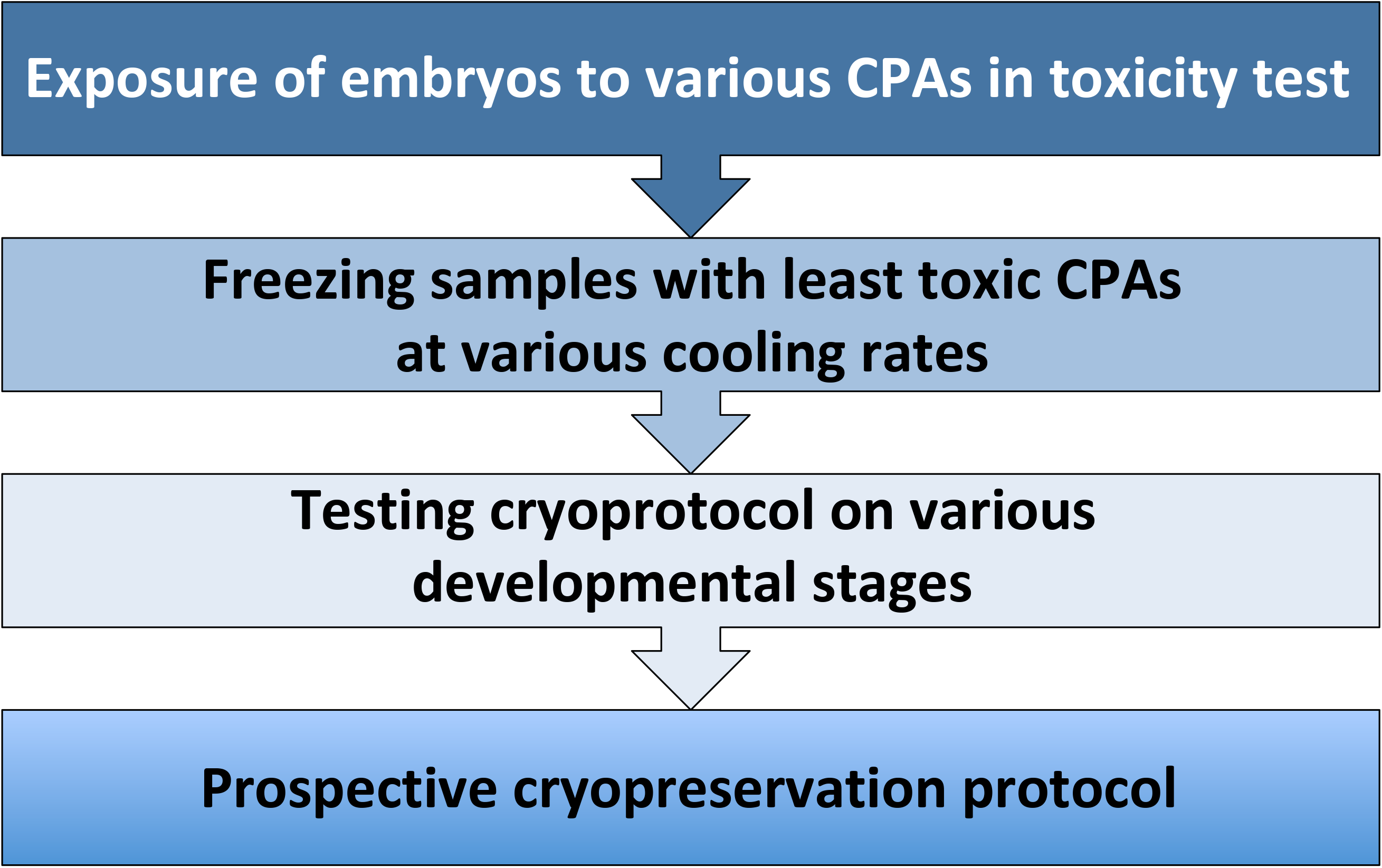
Flowchart illustrating the experimental workflow we used to identify an optimal cryopreservation process for *T. gratilla* embryos.

### Experiment #1: Toxicity Effects of Various Permeating and Non-permeating Cryoprotectants on Embryos

To establish an effective cryopreservation protocol for *T. gratilla*, urchin embryos in the gastrula stage were exposed to a variety of cryoprotectant agents in various concentrations, then their development was followed for three days after cryoprotectant agent exposure. This exploratory investigation employed cryoprotectant agents that were common in the literature, at various concentrations and combinations: dimethyl sulfoxide (1M and 2M; 1M Me2SO + 0.5M trehalose and 1M Me2SO + 0.125M trehalose), propylene glycol (1M and 2M), ethylene glycol (1M and 2M) and just trehalose (0.5M) [2,56,62].

For each treatment, approximately 100 embryos (116 ± 5) were aliquoted into 2-ml cryovials in 0.5 ml FSW to which 500 µl of test solutes were added. Rapid dehydration of embryos induced by cryoprotectant agent exposure can be lethal [44]. To diminish the damage incurred from potential osmotic stress, cryoprotectant agents were gradually introduced to the embryos in a stepwise fashion [2,56,62], by adding 50 µl of the test solutions in 10-equivolumetric steps (once/minute for 10 minutes) at room temperature, to a final volume of 1 ml. The larvae were then placed on ice and given 20 minutes to equilibrate.

After equilibration, the embryos were rehydrated at RT (23–25°C) in 10-equivolumetric steps (50 µl of FSW was added once/minute for 10 minutes). The embryos were then transferred to a 40-µm mesh filter basket to rinse in FSW before being transferred to a 35-mm petri dish and incubated at 27 °C. Embryonic development was assessed daily to compare the impacts of the different concentrations of assorted cryoprotectant agents against their untreated control counterparts.

Untreated controls were maintained in FSW at 27 °C in an incubator, with no exposure to cryoprotectant agents or chilling stress. All samples (around 100 embryos per replicate) were examined under a dissection microscope to estimate the proportion of embryo survival (as defined by motility and a clear appearance) as well as their respective stages of development. For every replicate from each treatment, embryos were counted to determine how many were still alive. Dead embryos tended to appear degraded and displayed no motility. For specimens that remained alive, each was categorized by its developmental stage (either gastrula, prism, 2-arm pluteus or 4-arm pluteus stages).

### Experiment #2: Testing Various Cryoprotectants and Cooling Rates

In order to devise a cryopreservation protocol, candidate cryoprotectants were used from the results of Experiment #1, specifically Me2SO, PG and EG, at a final concentration of 1M diluted in FSW (vol/vol). Approximately 100 embryos, in the gastrula stage, were aliquoted into 2-ml cryovials in 0.5 ml FSW. Candidate cryoprotectant agents were introduced to the embryos in 10-equivolumetric steps of 50 µL each (once/minute for 10 minutes) at room temperature, to a final volume of 1 ml in cryovials. The embryos were then placed on ice for 10 min to equilibrate, as recommended in the protocol by Odintsova et al. [52]. During this trial, different cooling rates (1.5°C/min, 4.7°C/min, 9.7°C/min and 15.2°C/min) were tested in a controlled-rate freezer (BioCool (FTS Systems Inc., Stone Ridge, NY, USA). These freezing rates were started at 0°C, until the samples hit –35°C, where they were held for 5 minutes, before being transferred and quenched in liquid nitrogen as in Odintsova et al. [52]. Cryovials were thawed by swirling in a heated water bath at 60°C for nearly 2 minutes until no visible ice was left in the vial, then rapidly returned to RT (23–25°C) prior to rehydration.

Rehydration and incubation protocols were then performed and embryonic development was monitored for three days after thawing, as described in Experiment #1 above.

In some cases, the addition of trehalose, a non-permeating cryoprotectant agent, has been documented to improve cryopreservation results by reducing the toxicity of Me2SO [56,8]. To ascertain if these benefits extend to the cryopreservation of *T. gratilla* larvae, the addition of trehalose (TRE) to the protocol was examined. Protocols with mixtures of 1M Me2SO with 0.1M TRE and 0.05M TRE were performed alongside treatments with 1M Me2SO alone for comparison. Embryos in the gastrula stage were transferred into cryovials to which cryoprotectant agents were introduced as described above. The samples were frozen at a cooling rate of -9.7°C/min to –35°C and held at that temperature for 5 minutes, before being transferred into liquid nitrogen. Thawing, rehydration, rinse out and incubation protocols were then performed as described above. Development of the embryos was assessed daily for three days after thawing.

### Experiment #3: Effect of Cryopreservation on Various Developmental Stages

Effectiveness of the cryopreservation protocol was assessed at four developmental stages. These were: 1) fertilized eggs (1 h post-fertilization); 2) blastula (10 h post-fertilization); 3) gastrula (18 h post-fertilization) and 4) 4-arm pluteus (48 h post-fertilization). Cryopreservation, thawing, rehydration and incubation protocols were conducted as described above in Experiment #2, using 1M Me2SO in FSW as the candidate cryoprotectant agent. Development and motility of the treated developmental stages were assessed daily for three days post-thaw and assessed as described in Experiment #1.

### Statistics

Data were analyzed and treatments were compared using Kaplan-Meier survivorship curves along with the non-parametric Wilcoxon test. Statistical tests were performed in RStudio (RStudio, Inc.) [61], using the “survival” [71], “survminer” [39], with “ggplot2” [75] packages and JMP Pro 11 (SAS Institute, Inc.).

## Results

### Experiment #1: Toxicity

Examination of *T. gratilla* gastrulae subjected to toxicity trials revealed a broad range of survival. In general, embryos exposed to higher concentrations of cryoprotectant agents (such as 2M EG and 2M PG) suffered lower survival over the course of the experiment (2-39%) (Figure S1 & S2) (*p* < 0.05, Wilcoxin pairwise comparison) (Table S1-A). These low survival rates were not found to be different from treatments exposed to 2M Me2SO (*p* > 0.05, Wilcoxin pairwise comparison) (Table S1-A), which also had a broad range of survival rates (50.5 ± 28.6%).

Embryos exposed to moderate concentrations of cryoprotectant agents (1M Me2SO, 1M PG and 1M EG, 1M Me2SO + 0.125M TRE, 0.5M TRE, 1M Me2SO + 0.5M TRE) experienced the highest survival rates (69.8-100%), of which treatments using 1M Me2SO had the highest survival (100%) followed by treatments using 1M Me2SO + 0.125M TRE (96.1 ± 3.1%) then 1M EG (95.2 ± 4.8%)and 1M PG (86 ± 14.5%), followed by 0.5M TRE (76.4 ± 8.5%) and then 1M Me2SO + 0.05M TRE (69.8 ± 17.9%). The percent survival of these treatments was statistically indistinguishable from each other or to the control treatments, which were not exposed to any cryoprotectant agents and had 100% survival (*p* > 0.05, Wilcoxin pairwise comparison) (Table S1-A) (Figure S2).

When looking beyond survival, and considering the development of embryos to the most advanced, observed, development stage (4-arm pluteus larvae) across treatments, a different pattern emerged. In the control treatment, 96.5 ± 1.4% of embryos developed to the 4-arm pluteus stage (Figure 2). By the third day, 91.7 ± 4.9% of embryos treated with 1M EG developed to the 4-arm pluteus stage, 67.7 ± 23.1% of embryos developed to the 4-arm pluteus stage from the 1M PG treatments, and 55.8 ± 23.9% of embryos treated with 1M Me2SO developed to the 4-arm pluteus stage (Figure 2). Treatments exposing embryos to 1M EG, 1M PG and 1M Me2SO were not found to be statistically distinguishable from the control (*p* > 0.05, Wilcoxin pairwise comparison) (Figure 2) (Table S1-B). Rates of development to the 4-arm pluteus stage were lower than the controls (∼97%), for the other tested treatments, with only 25.1 ± 14.6% for 0.5M TRE, 16.2 ± 15.7% for 1M Me2SO + 0.125M, 14.4 ± 11.9% for 2M Me2SO, 12.4 ± 1.6% for 2M EG, 2.8 ± 2.8% for 1M Me2SO + 0.5M TRE and 1.3 ± 1.0% for 2M PG (*p* < 0.05 Wilcoxin pairwise comparison) (Figure 2) (Table S1-B).

**Figure 2:**
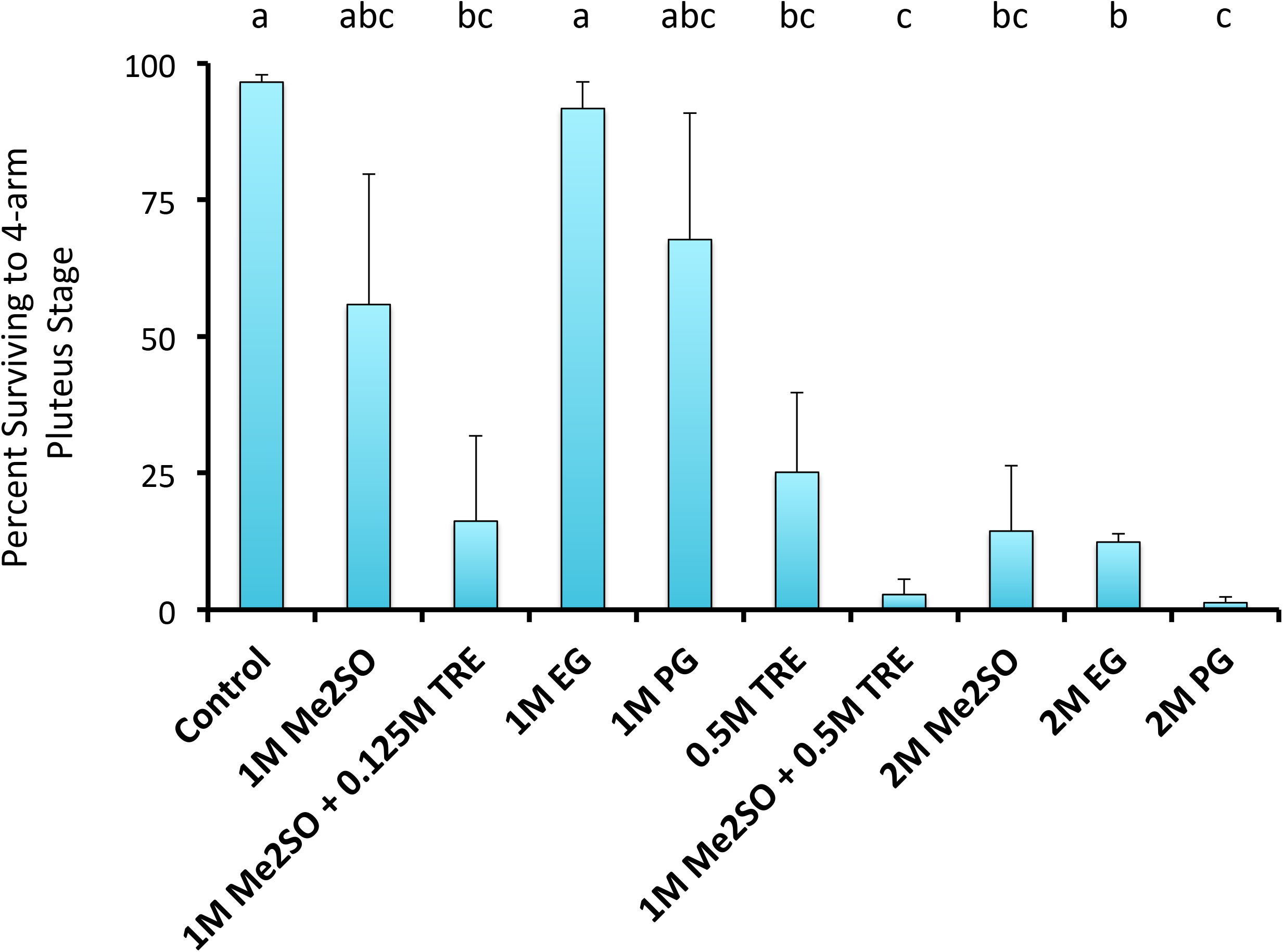
Proportion of *Tripneustes gratilla* offspring that reached the 4-arm pluteus stage of development 72 hours post exposure to various concentrations and mixtures of cryoprotectant agents (Me2SO = dimethyl sulfoxide, PG = propylene glycol, EG = ethylene glycol, TRE = trehalose). Cryoprotectant exposure occurred during gastrula stage. Note letters identify significant subsets (*p* < 0.05 Wilcoxin pairwise comparison) (n=6 females; 3 pooled males and 100 larvae/treatment).

### Experiments #2: Cryopreservation Experiments—Testing cryoprotectant agents and cooling rates

When analyzing the variability in survival of urchin embryos cryopreserved with different cryoprotectant agents (1M Me2SO, 1M PG and 1M EG) and at different cooling rates (-15.2°C/min, -9.7°C/min, -4.7°C/min and -1.5°C/min), only a few of the tested protocols yielded viable offspring (Figure S3). Embryonic survival rates between protocols were found to be statistically different (*p* < 0.05, Kaplan-Meier survivorship curves, Figure S3). None of the tested cryoprotectant agents produced any surviving embryos when samples were cooled at a rate of -1.5°C/min (Figure S3 & S4). Treatments using a cooling rate of -4.7°C/min had a 0% survival rate for all cryoprotectant agents, except PG, which had a survival rate of 34.4 ± 9.2% (Figure S3 & S4). Protocols with a cooling rate of -9.7°C/min produced viable offspring with 1M Me2SO and 1M EG (71.4 ± 3.9% and 23.2 ± 4.1%, respectively), but none survived at this cooling rate when using PG as cryoprotectant (Figure S3 & S4). Treatments using a cooling rate of -15.2°C/min produced survivors only with 1M Me2SO and 1M EG (37.6 ± 6.8% and 14.8 ± 5.0%, respectively) (Figure S3 & S4). *Tripneustes gratilla* embryos treated with 1M Me2SO and frozen with a cooling rate of -9.7°C/min had statistically higher survival than embryos treated with all other tested cryopreservation protocols (Figure S4) (*p* < 0.05, Wilcoxin pairwise comparison) (Table S2-A). Although the survival of treatments cryopreserved with 1M Me2SO cooled at -9.7°C/min were slightly less than the FSW controls (71.3±3.9% vs. 76.1±8.6%, respectively), these differences were found to be statistically indecipherable (*p* = 0.7469, Wilcoxon pairwise comparison) (Figure S3, S4 & Table S2-A).

A similar pattern emerges when analyzing just the data of cryopreserved embryos that continued their development to the 4-arm pluteus stage. Of the samples that were cryopreserved, treatments cooled at -9.7°C/min and -15.2°C/min with 1M Me2SO produced the highest proportion of embryos that developed to the 4-arm pluteus stage (25.9±7.2% and 10.4±4.2%, respectively), which were not statistically distinguishable from each other (Figure 3)(*p* = 0.1146, Wilcoxon pairwise comparison) (Table S2-B); however, samples treated with 1M Me2SO and cooled at -9.7°C/min did produce more larvae at the advanced 4-arm pluteus stage than all other treatments. The only other treatments that produced 4-arm pluteus stage larvae were those cryopreserved with 1M PG at -4.7°C/min and 1M EG at -9.7°C/min (6.9±3.5% vs. 1.6±0.7%, respectively) (Figure 3). No other cryopreservation treatments produced 4-arm pluteus stage larvae. Also, samples cooled at -9.7°C/min with 1M Me2SO produced high enough proportions of 4-arm pluteus stage larvae that this treatment was not found to be statistically different from the control) (*p* > 0.05, Wilcoxon pairwise comparison) (Table S2-B).

**Figure 3:**
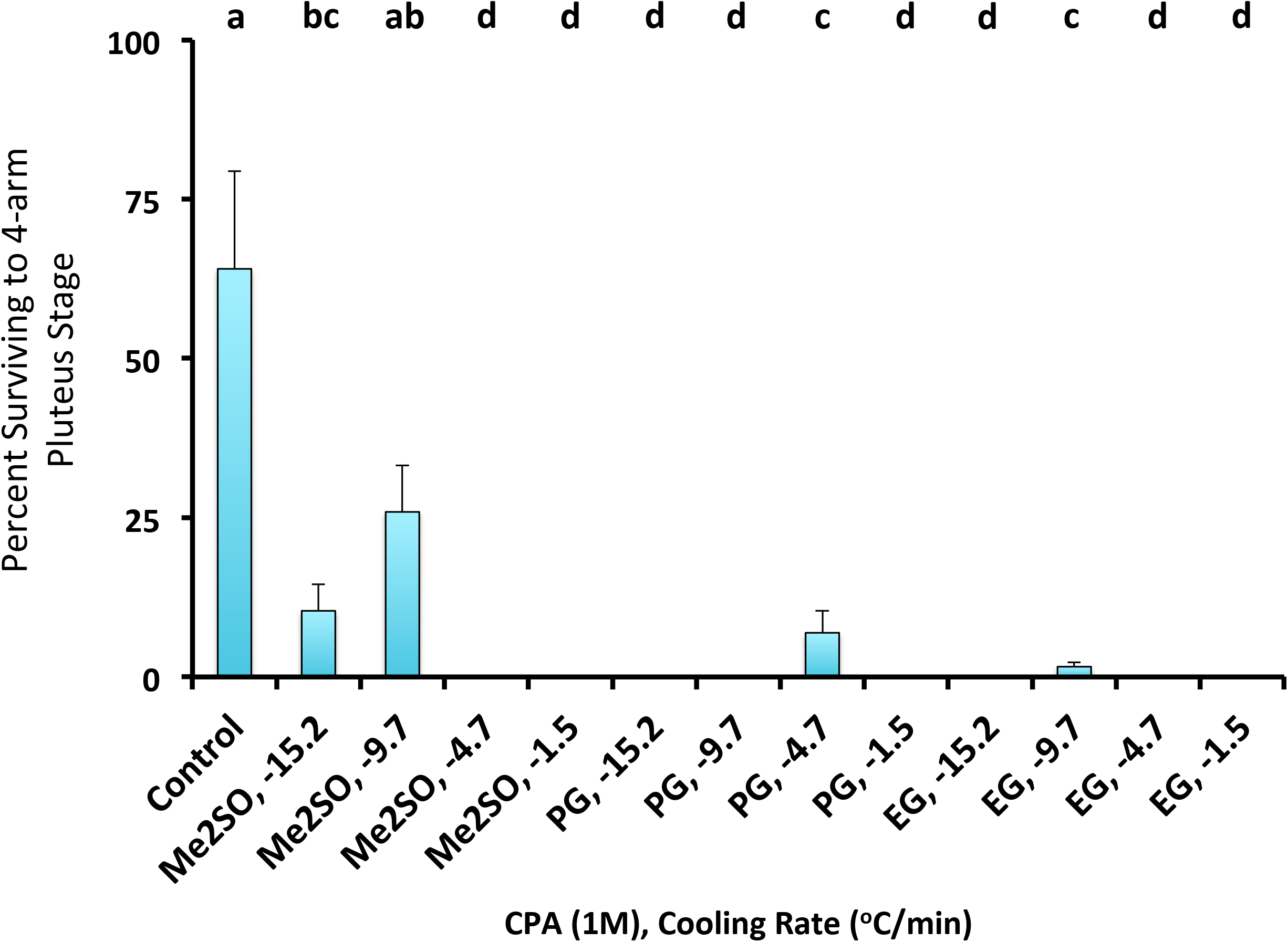
Proportion of *Tripneustes gratilla* offspring, tested at the gastrula stage, that developed to the 4-arm pluteus stage 72 hours post thawing from cryofreezing at different cooling rates (-15.2°C/min, -9.7°C/min, -4.7°C/min and -1.5°C/min) and with various cryoprotectant agents (n=8). Abbreviations: Me2SO = dimethyl sulfoxide, PG = propylene glycol, EG = ethylene glycol. Note letters identify significant subsets (*p* < 0.05 Wilcoxin pairwise comparison).

### Experiment #3: Effect of Cryopreservation on Various Developmental Stages

*Tripneustes gratilla* offspring at various early stages of ontogenetic development were exposed to the cryopreservation protocol to compare their performance post-thaw. The differences in survival rates between cryopreserved stages were found to be different (*p* < 0.05, Kaplan-Meier survivorship curves, Figure S7). No survival was achieved while attempting to cryopreserve fertilized eggs (Figure S7 & S8) (*p* < 0.05 Wilcoxin pairwise comparison) (Table S4). Larvae cryopreserved at the 4-arm pluteus stages experienced the next lowest survival, with only 27 ± 2.8% remaining 72 h post-thaw (Figure 4, S7 & S8). Embryos cryopreserved at the blastula stage had intermediate survivorship, with 48.3 ± 7.8% remaining 72 h post-thaw, and although embryos cryopreserved in the blastula stage had higher survival than 4-arm pluteus stage larvae (48.2 ± 7.8% and 27.0 ± 2.8%, respectively), these differences were not found to be statistically distinct (Figure S7 & S8) (*p* = 0.066 Wilcoxin pairwise comparison) (Table S4).

**Figure 4:**
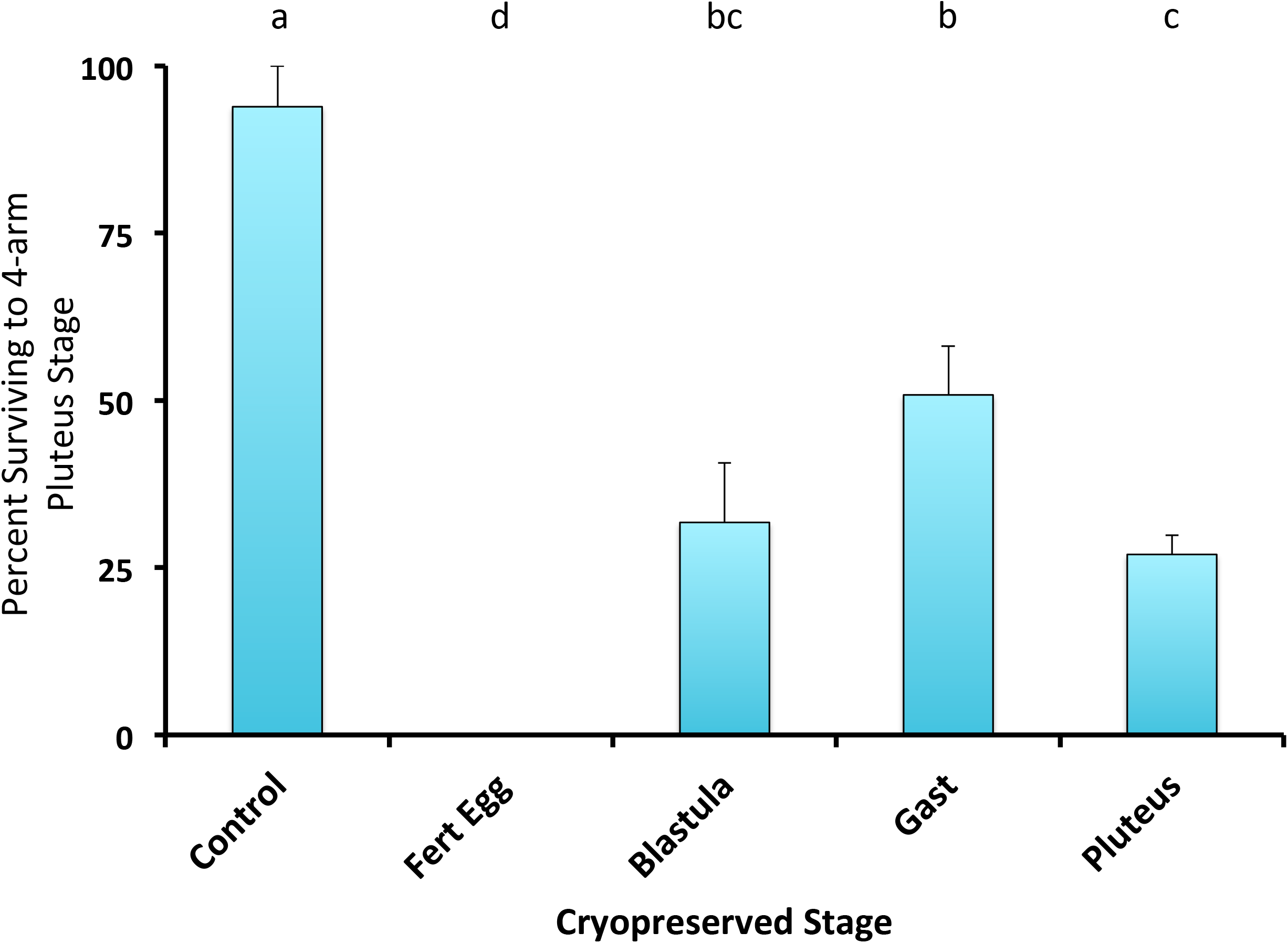
Proportion of *Tripneustes gratilla* offspring cryopreserved (with 1M Me2SO at -9.7°C/min) at various stages of development (fertilized egg, blastula, gastrula and pluteus) that continued to develop to the 4-arm pluteus stage, 72 h post-thaw (n=8). Note letters identify significant subsets (*p* < 0.05 Wilcoxin pairwise comparison).

A similar pattern was observed when comparing the proportion of blastulae that developed to the 4-arm pluteus stage, 31.8 ± 8.9% (Figure 4). Embryos cryopreserved in the gastrula stage exhibited higher survivorship rates (77.5 ± 6.7%) than fertilized eggs and 4-arm pluteus stage larvae (0% and 27 ± 2.8%, respectively) (*p* < 0.05, Wilcoxin pairwise comparison), and marginally higher survival (48.3 ± 7.8%) than treatments cryopreserved in the blastula stage (*p* = 0.052, Wilcoxin pairwise comparison) (Figure S7 & S8) (Table S4). A similar pattern emerges when comparing development rates; 50.8 ± 7.3% of cryopreserved gastrulae matured to the 4-arm pluteus stage, which was found to be statistically higher than the survival of larvae that were already at the 4-arm pluteus stage at the time of cryopreservation (Figure 4, *p* = 0.041, Wilcoxin pairwise comparison, Table S5). Although cryopreserved gastrulae appeared to have higher maturation rates when compared to the cryopreserved blastulae (31.8 ± 8.9% developed to the 4-arm pluteus stage), these two treatments were not different (Figure 4, *p* > 0.05, Wilcoxin pairwise comparison)(Table S5).

## Discussion

This study aimed to extend the benefits of artificial reproductive techniques through cryopreservation to the collector urchin, *T. gratilla*. Although certain urchins, such as *Evechinus chloroticus*, have exhibited favorable responses to high (2M) concentrations of permeating cryoprotectant agents [2], our experiments indicate that *T. gratilla* embryos exhibit poor survival and development when exposed to 2M concentrations of cryoprotectant agents, a pattern analogous to the urchin *Paracentrotus lividus* [56]. Several have reported on the benefits of non-permeating cryoprotectant agents (such as trehalose) when cryopreserving urchin embryos [8, 52], yet these benefits do not extend to all urchin species [2]. Trends in our data suggest that embryonic development is stunted by the addition of the non-permeant cryoprotectant agent trehalose, whether the samples are cryopreserved or just exposed to the agent.

Slow cooling rates (1-2.5°C/min) have been effective for the cryopreservation of embryos of other urchin species (such as *P. lividus* and *E. chloroticus*) [2,52, 56], yet slow cooling (1.5°C/min) was least effective with *T. gratilla* embryos, causing 0% survival with all tested cryoprotectant agents. However, not all urchin embryos need to be cooled at slow rates to achieve high survival, as *Strongylocentrotus intermedius* has been demonstrated to tolerate a wide range of cooling rates (4-40°C/min) [5]. Examining an assortment of cooling regimes revealed that chilling rates ranging from ∼5°C/min to ∼15°C/min could produce surviving larvae when using 1M concentrations of either Me2SO, EG or PG for *T. gratilla*. Embryos can exhibit different levels of permeability from one cryoprotectant agent to the next, potentially leading to observed variation in cryopreservation efficiency [29]. For instance, embryos exposed to EG in our toxicity trials exhibited high survival and maturation rates, however embryos performed poorly at all observed cooling rates when cryopreserve with this cryoprotectant agent, potentially indicating that EG did not effectively permeate into *T. gratilla* cells.

Although some have managed to document growth of fertilized eggs after cryopreservation [56], no successes have been documented for oocyte cryopreservation in urchins, or other marine organisms [57,14], and no survival was achieved while attempting to cryopreserve fertilized eggs in *T. gratilla* in the present study. Eggs are inherently more difficult to cryopreserve because of their membranes’ poor permeability to water and cryoprotective solutes, both of which are indispensable to successful cryopreservation [48]. Evaluating the survival and maturation of various life stages, post freezing, established that *T. gratilla* can be cryopreserved at the blastula, gastrula and 4-arm echinopluteus stages, albeit with variable success. Similarly, cryopreservation protocols developed for other urchin species have reported disparate performances between embryonic and larval stages. For example, *Hemicentrotus pulcherrimus,E. chloroticus*, *S. intermedius* and *S. nudus* showed the most promise when cryopreserved as 4-arm pluteus larvae [2, 4, 5], while the highest performance for *P. lividus* was reported when cryopreserving blastulae [56]. Such variable cryopreservation efficiency can occur from changes in membrane permeability and surface-to-volume ratios of cells while an organism develops [4, 30]. For *T. gratilla*, blastulae and gastrulae developmental stages may benefit from a lack of rigid body parts, allowing them to “bounce back” from the squeezing forces of extracellular ice. As embryonic development proceeds, the invagination of the archenteron would increase the cells’ surface area, conceivably increasing transportation of water and cryoprotectant agents across membranes, potentially accounting for the improved performance of cryopreserved *T. gratilla* gastrulae.

In our tests, collector urchins exhibited different levels of sensitivity to all examined cryopreservation protocols, with many being completely fatal to our specimens. Nevertheless, a promising cryopreservation protocol for *T. gratilla* was elucidated during this study. Using 1M Me2SO on the gastrula stage with a cooling rate of ∼10°C/min consistently produced samples with the highest survival and maturation rates. In some assessments, our cryopreservation protocol produced offspring with survival that were statistically indistinguishable from our control treatments, suggesting that cryopreservation may be carried out with negligible impacts to the survivorship of *T. gratilla*’s early life stages. Future directions for this study will include the set up of grow out cultures in order to assess the performance of cryopreserved embryos throughout ontogenetic development. Fine-tuning parameters of our established protocol could optimize survival and development of cryopreserved *T. gratilla*, and rearing cryopreserved larvae to the point of competency and settlement would provide a proof of concept that this protocol could be applied to the aquaculture of this species.

Collector urchins play a valuable role on Hawaiian reefs by contributing to the delicate balance between algae and coral growth. Introduced algae species, which turned invasive, have plagued Kane’ohe Bay’s reefs since the 1970s, and threatened to proliferate around Oahu if not contained [63]. Due to the algae’s fast recovery after manual removal efforts, investigation into potential biocontrol agents led researchers to promoting *T. gratilla*, in concert with manual removal [66,20]. As such, the state has spent considerable time and resources developing and maintaining an aquaculture facility dedicated to producing urchins for use as biocontrol. Cryopreservation of reproductive materials during peak breeding seasons could become a systematic approach when harvesting urchins for aquaculture, in the same way that banking has for some corals [28]. Establishing a frozen biorepository (as has been done for corals [33, 35] could contribute to more robust management strategies to aid in reef restoration efforts. As a conservation tool for coral reefs, cryopreservation has potential to preserve genetic diversity, supplement shrinking populations that suffer from losses in heterozygosity and genetic drift, as well as sustaining biodiversity by preventing extinctions [32]. Maintaining a consistent collection of contemporary samples would ensure frozen specimens from the repository did not become genetically and adaptively obsolete, in relation to wild populations of conspecifics. Thus, investigating the challenges of scaling-up these laboratory protocols will be critical in producing high throughput methods for *T. gratilla* aquaculture [27].

Continued breakthroughs in artificial reproduction, such as cryopreservation, along with the up scaling of these methods could also have applications to commercial aquaculture of *T. gratilla*. Due to the high demand for sea urchins as culinary delicacies, wild stocks have been overfished to the point of worldwide decline [65]. Overfishing of the congeneric *T. ventricosus* has caused the collapse of the sea urchin fishery and subsequent disappearance from the reef community in Barbados [64]. A similar local disappearance of *T. gratilla*, attributed to overexploitation, was documented in 1992 in the Philippines [38]. Large-scale *T. gratilla* fisheries, responsible for catching hundreds of tons of urchins, have been reported in the Philippines, Fiji and Japan [3]. Urchin gonads, known as roe, are a luxury delicacy that can command astronomical prices, over US$1000/kg, making them one of the most expensive seafood products in the world [11,58]. With the ever-increasing demand for roe and the general decline in sea urchin fisheries, an opportunity for aquaculture and stock enhancement is bourgeoning [40,65]. Groups in Ecuador and Australia have developed and expanded aquaculture protocols geared at commercial production of *Tripneustes* urchins for the seafood market [51,67]. To assuage the pressure put on overexploited urchins, supplementation of wild populations has been proposed through the release of cultured specimens [38,25]. In Japan and the Philippines reseeding efforts have been established for over 30 years, in an attempt to compensate for the urchins removed from the reef by fisheries [3].

Coral reefs are being severely impacted by climate change [40], causing parallel impacts on urchin populations, health and viability due to stress and disease. Urchins are critical for maintaining the balance of reefs so that they do not shift into an algal-dominated state. This may be the first time that a tropical sea urchin has been cryopreserved, as most protocols developed for urchins have focused on temperate species. While work continues on *T. gratilla*, it is hoped that these techniques can be extended and adapted to other tropical urchin species. Artificial reproductive techniques could become increasingly important for reseeding reefs where ecologically vital urchins have become extirpated, such as *Diadema antillarum* in the Caribbean. The cryopreservation of these important tropical urchin species will be and essential tool for reef restoration in the future.

## Acknowledgements

Thank you to Mariko Westbrook, Brock Ball, Mason Barca and Evan Barba for their assistance specimen collections. This is contribution XXXX from the Hawaìi Institute of Marine Biology and XXXX from the School of Ocean and Earth Science and Technology.

## Authors’ contributions

**Conceived and designed the analysis: CW, MH, JD**

**Collected the data: CW**

**Contributed data or analysis tools: CW, MH**

**Performed the analysis: CW**

**Wrote the paper: CW, BB, MH, JD.**

## Accompanying Legends

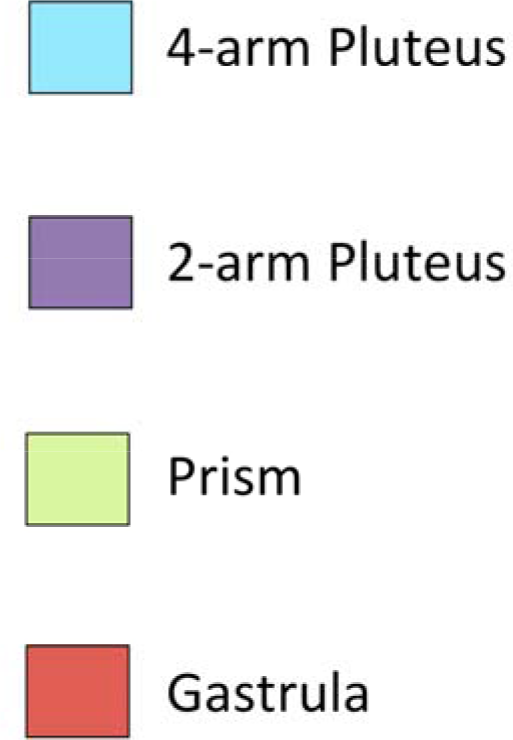
Legend for Figure S2, S4, S6 and S8

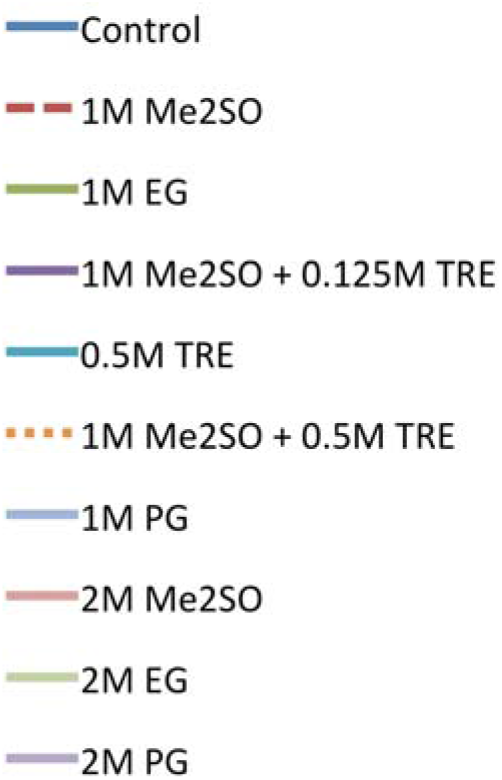
Legend for Figure S1

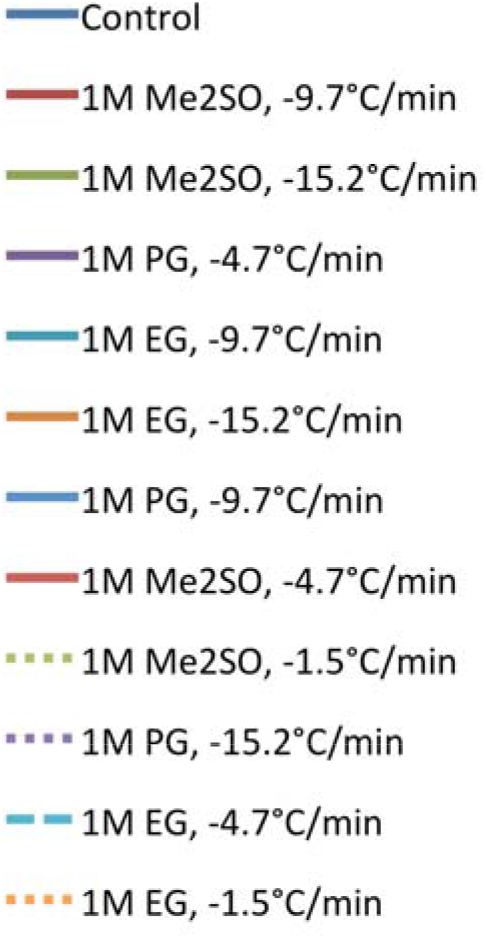
Legend for Figure S3

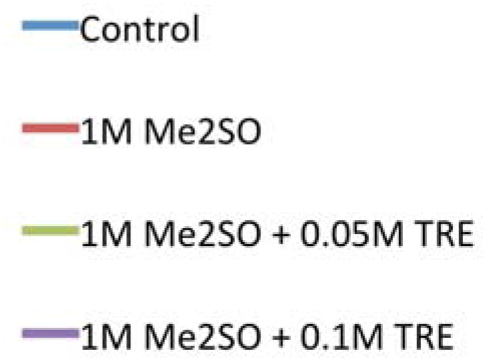
Legend for Figure S5

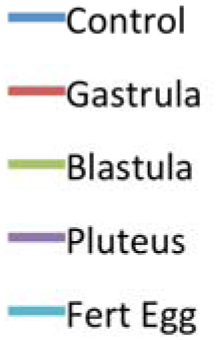
Legend for Figure S7

