## Supplemental Methods and Data for "Cryopreservation of the collector urchin embryo, *Tripneustes gratilla*"

1) Toxicity Experiment

Figure S1: Kaplan-Meier product-limit survivorship curves of toxicity trials for *Tripneustes gratilla* embryos exposed to a variety of cryoprotectant agents at the gastrula stage (n=600).

Abbreviations: TRE = trehalose, Me2SO = dimethyl sulfoxide, EG = ethylene glycol, PG = propylene glycol, FSW = filtered seawater.

Table S1-A: Comparison of survivorship rates between toxicity trial treatments for *Tripneustes gratilla*. Abbreviations: TRE = trehalose, Me2SO = dimethyl sulfoxide, EG = ethylene glycol, PG = propylene glycol, FSW = filtered seawater.

| **Between** | **df** | **Wilcoxin** | | **Survivorship Verdict** |
| --- | --- | --- | --- | --- |
|  |  | **Statistic** | **P** |  |
| All treatments | 9 | 25.2927 | 0.0027 |  |
| 0.5M TRE vs 1M Me2SO | 1 | 1.81895 | 0.0689 | NOT SIGNIFICANT |
| 0.5M TRE vs 1M Me2SO + 0.125M TRE | 1 | 1.33112 | 0.1832 | NOT SIGNIFICANT |
| 0.5M TRE vs 1M Me2SO + 0.5M TRE | 1 | -0.14790 | 0.8824 | NOT SIGNIFICANT |
| 0.5M TRE vs 1M EG | 1 | 0.97125 | 0.3314 | NOT SIGNIFICANT |
| 0.5M TRE vs 1M PG | 1 | 0.61512 | 0.5385 | NOT SIGNIFICANT |
| 0.5M TRE vs 2M Me2SO | 1 | -0.14790 | 0.8824 | NOT SIGNIFICANT |
| 0.5M TRE vs 2M EG | 1 | -2.45181 | 0.0142 | 0.5M TRE > 2M EG |
| 0.5M TRE vs 2M PG | 1 | -2.17807 | 0.0294 | 0.5M TRE > 2M PG |
| 0.5M TRE vs FSW | 1 | 1.81895 | 0.0689 | NOT SIGNIFICANT |
| 1M Me2SO vs 1M Me2SO + 0.125M TRE | 1 | -1.32288 | 0.1859 | NOT SIGNIFICANT |
| 1M Me2SO vs 1M Me2SO + 0.5M TRE | 1 | -1.32288 | 0.1859 | NOT SIGNIFICANT |
| 1M Me2SO vs 1M EG | 1 | -1.80144 | 0.0716 | NOT SIGNIFICANT |
| 1M Me2SO vs 1M PG | 1 | -0.75000 | 0.4533 | NOT SIGNIFICANT |
| 1M Me2SO vs 2M Me2SO | 1 | -1.32288 | 0.1859 | NOT SIGNIFICANT |
| 1M Me2SO vs 2M EG | 1 | -2.52966 | 0.0114 | 1M Me2SO > 2M EG |
| 1M Me2SO vs 2M PG | 1 | -2.32246 | 0.0202 | 1M Me2SO > 2M PG |
| 1M Me2SO VS FSW | 1 | 0 | 1 | NOT SIGNIFICANT |
| 1M Me2SO + 0.125M TRE vs 1M Me2SO + 0.5M TRE | 1 | -0.46134 | 0.6446 | NOT SIGNIFICANT |
| 1M Me2SO + 0.125M TRE vs 1M EG | 1 | 0.00000 | 1.0000 | NOT SIGNIFICANT |
| 1M Me2SO + 0.125M TRE vs 1M PG | 1 | 0.16536 | 0.8687 | NOT SIGNIFICANT |
| 1M Me2SO + 0.125M TRE vs 2M Me2SO | 1 | -0.46134 | 0.6446 | NOT SIGNIFICANT |
| 1M Me2SO + 0.125M TRE vs 2M EG | 1 | -2.45927 | 0.0139 | 1M Me2SO + 0.125M TRE > 2M EG |
| 1M Me2SO + 0.125M TRE vs 2M PG | 1 | -2.19131 | 0.0284 | 1M Me2SO + 0.125M TRE > 2M PG |
| 1M Me2SO + 0.125M TRE vs FSW | 1 | 1.32288 | 0.1859 | NOT SIGNIFICANT |
| 1M Me2SO + 0.5M TRE vs 1M EG | 1 | 0.32996 | 0.7414 | NOT SIGNIFICANT |
| 1M Me2SO + 0.5M TRE vs 1M PG | 1 | 0.49608 | 0.6198 | NOT SIGNIFICANT |
| 1M Me2SO + 0.5M TRE vs 2M Me2SO | 1 | -0.46134 | 0.6446 | NOT SIGNIFICANT |
| 1M Me2SO + 0.5M TRE vs 2M EG | 1 | -1.39002 | 0.1645 | NOT SIGNIFICANT |
| 1M Me2SO + 0.5M TRE vs 2M PG | 1 | -2.19131 | 0.0284 | 1M Me2SO + 0.5M TRE > 2M PG |
| 1M Me2SO + 0.5M TRE vs FSW | 1 | 1.32288 | 0.1859 | NOT SIGNIFICANT |
| 1M EG vs 1M PG | 1 | 0.56857 | 0.5696 | NOT SIGNIFICANT |
| 1M EG vs 2M Me2SO | 1 | -0.32996 | 0.7414 | NOT SIGNIFICANT |
| 1M EG vs 2M EG | 1 | -2.80715 | 0.0050 | 1M EG > 2M EG |
| 1M EG vs 2M PG | 1 | -2.46680 | 0.0136 | 1M EG > 2M PG |
| 1M EG vs FSW | 1 | 1.80144 | 0.0716 | NOT SIGNIFICANT |
| 1M PG vs 2M Me2SO | 1 | -0.82680 | 0.4084 | NOT SIGNIFICANT |
| 1M PG vs 2M EG | 1 | -1.83458 | 0.0666 | NOT SIGNIFICANT |
| 1M PG vs 2M PG | 1 | -2.23253 | 0.0256 | 1M PG > 2M PG |
| 1M PG vs FSW | 1 | 0.75000 | 0.4533 | NOT SIGNIFICANT |
| 2M Me2SO vs 2M EG | 1 | 0.00000 | 1.0000 | NOT SIGNIFICANT |
| 2M Me2SO vs 2M PG | 1 | -1.04185 | 0.2975 | NOT SIGNIFICANT |
| 2M Me2SO vs FSW | 1 | 1.32288 | 0.1859 | NOT SIGNIFICANT |
| 2M EG vs 2M PG | 1 | -2.45927 | 0.0139 | 2M EG > 2M PG |
| 2M EG vs FSW | 1 | 2.52966 | 0.0114 | 2M EG < FSW |
| 2M PG vs FSW | 1 | 2.32246 | 0.0202 | 2M PG < FSW |

Figure S2: Proportion of *Tripneustes gratilla* offspring at various stages of development (gastrula, prism, 2-arm pluteus, 4-arm pluteus, as well as dead) 72 h post-exposure to various concentrations and mixtures cryoprotectant agents (Me2SO = dimethyl sulfoxide, PG = propylene glycol, EG = ethylene glycol, TRE = trehalose). Cryoprotectant exposure occurred during embryonic stage. Note letters identify significant subsets (*p* < 0.05 Wilcoxin pairwise comparison) (n=6).

2-A) Testing Various Cryoprotectant Agents and Cooling Rates

Figure S3: Kaplan-Meier product-limit survivorship curves of *Tripneustes gratilla* embryo cryopreserved at the gastrula stage with different cryoprotectant agents, and at different cooling rates (-15.2^o^C/min, -9.7^o^C/min, -4.7^o^C/min and -1.5^o^C/min) (n=800). Abbreviations: Me2SO = dimethyl sulfoxide, PG = propylene glycol, EG = ethylene glycol, FSW = filtered seawater.

Table S1-B: Comparison of larval development rates to the 4-arm pluteus stage between toxicity trial treatments for *Tripneustes gratilla*. Abbreviations: TRE = trehalose, Me2SO = dimethyl sulfoxide, EG = ethylene glycol, PG = propylene glycol, FSW = filtered seawater.

| **Between** | **df** | **Wilcoxin** | | **Survivorship Verdict** |
| --- | --- | --- | --- | --- |
|  |  | **Statistic** | **P** |  |
| All treatments | 9 | 25.6166 | 0.0024 |  |
| 1M EG vs 0.5M TRE | 1 | 2.45927 | 0.0139 | 1M EG > 0.5M TRE |
| 1M EG vs 1M Me2SO + 0.125M TRE | 1 | 2.45927 | 0.0139 | 1M EG > 1M Me2SO + 0.125M TRE |
| 1M EG vs 1M Me2SO + 0.5M TRE | 1 | 2.48208 | 0.0139 | 1M EG > 1M Me2SO + 0.5M TRE |
| FSW vs 2M EG | 1 | 2.45927 | 0.0139 | FSW > 2M EG |
| 2M EG vs 1M Me2SO + 0.5M TRE | 1 | 2.05041 | 0.0403 | 2M EG > 1M Me2SO + 0.5M TRE |
| FSW vs 0.5M TRE | 1 | 2.19131 | 0.0284 | FSW > 0.5M TRE |
| FSW vs 1M Me2SO + 0.125M TRE | 1 | 2.19131 | 0.0284 | FSW > 1M Me2SO + 0.125M TRE |
| FSW vs 1M Me2SO + 0.5M TRE | 1 | 2.23253 | 0.0256 | FSW > 1M Me2SO + 0.5M TRE |
| FSW vs 2M Me2SO | 1 | 2.17807 | 0.0294 | FSW > 2M Me2SO |
| FSW vs 2M PG | 1 | 2.19131 | 0.0284 | FSW > 2M PG |
| 1M PG vs 1M Me2SO + 0.5M TRE | 1 | 1.53781 | 0.1241 | NOT SIGNIFICANT |
| 2M EG vs 1M Me2SO + 0.125M TRE | 1 | 1.17617 | 0.2395 | NOT SIGNIFICANT |
| 2M EG vs 2M Me2SO | 1 | 1.17260 | 0.2410 | NOT SIGNIFICANT |
| 1M PG vs 0.5M TRE | 1 | 1.33112 | 0.1832 | NOT SIGNIFICANT |
| 1M PG vs 1M Me2SO + 0.125M TRE | 1 | 1.33112 | 0.1832 | NOT SIGNIFICANT |
| 1M EG vs 1M Me2SO | 1 | 0.95940 | 0.3374 | NOT SIGNIFICANT |
| 1M Me2SO VS FSW | 1 | 1.01643 | 0.3094 | NOT SIGNIFICANT |
| 2M Me2SO vs 1M Me2SO + 0.5M TRE | 1 | 0.92269 | 0.3562 | NOT SIGNIFICANT |
| 1M Me2SO vs 0.5M TRE | 1 | 0.73951 | 0.4596 | NOT SIGNIFICANT |
| FSW vs 1M PG | 1 | 0.72602 | 0.4678 | NOT SIGNIFICANT |
| 2M Me2SO vs 1M Me2SO + 0.125M TRE | 1 | 0.44371 | 0.6573 | NOT SIGNIFICANT |
| 1M PG vs 1M Me2SO | 1 | 0.14609 | 0.8839 | NOT SIGNIFICANT |
| 2M PG vs 1M Me2SO + 0.5M TRE | 1 | 0.16536 | 0.8687 | NOT SIGNIFICANT |
| FSW vs 1M EG | 1 | 0.10692 | 0.9148 | NOT SIGNIFICANT |
| 1M Me2SO + 0.125M TRE vs 0.5M TRE | 1 | 0.00000 | 1.0000 | NOT SIGNIFICANT |
| 2M Me2SO vs 0.5M TRE | 1 | 0.00000 | 1.0000 | NOT SIGNIFICANT |
| 2M EG vs 0.5M TRE | 1 | 0.00000 | 1.0000 | NOT SIGNIFICANT |
| 2M PG vs 1M Me2SO + 0.125M TRE | 1 | -0.15378 | 0.8778 | NOT SIGNIFICANT |
| 1M Me2SO + 0.5M TRE vs 1M Me2SO + 0.125M TRE | 1 | -0.49608 | 0.6198 | NOT SIGNIFICANT |
| 2M PG vs 0.5M TRE | 1 | -0.46134 | 0.6446 | NOT SIGNIFICANT |
| 1M PG vs 1M EG | 1 | -0.53300 | 0.5940 | NOT SIGNIFICANT |
| 1M Me2SO + 0.5M TRE vs 0.5M TRE | 1 | -0.82680 | 0.4084 | NOT SIGNIFICANT |
| 2M Me2SO vs 1M Me2SO | 1 | -0.87123 | 0.3836 | NOT SIGNIFICANT |
| 1M Me2SO + 0.125M TRE vs 1M Me2SO | 1 | -1.03531 | 0.3005 | NOT SIGNIFICANT |
| 2M PG vs 2M Me2SO | 1 | -1.03531 | 0.3005 | NOT SIGNIFICANT |
| 2M Me2SO vs 1M PG | 1 | -1.16164 | 0.2454 | NOT SIGNIFICANT |
| 2M PG vs 1M Me2SO | 1 | -1.33112 | 0.1832 | NOT SIGNIFICANT |
| 2M PG vs 1M PG | 1 | -1.33112 | 0.1832 | NOT SIGNIFICANT |
| 2M EG vs 1M Me2SO | 1 | -1.17260 | 0.2410 | NOT SIGNIFICANT |
| 2M EG vs 1M PG | 1 | -1.17260 | 0.2410 | NOT SIGNIFICANT |
| 1M Me2SO + 0.5M TRE vs 1M Me2SO | 1 | -1.53781 | 0.1241 | NOT SIGNIFICANT |
| 2M Me2SO vs 1M EG | 1 | -2.45181 | 0.0142 | 2M Me2SO < 1M EG |
| 2M PG vs 1M EG | 1 | -2.45927 | 0.0139 | 2M PG < 1M EG |
| 2M PG vs 2M EG | 1 | -2.45927 | 0.0139 | 2M PG < 2M EG |
| 2M EG vs 1M EG | 1 | -2.80224 | 0.0051 | 2M EG < 1M EG |

Figure S4: Proportion of *Tripneustes gratilla* offspring at various stages of development (gastrula, prism, 2-arm pluteus, 4-arm pluteus) 72 hours post thawing from cryopreservation at different cooling rates (-15.2^o^C/min, -9.7^o^C/min, -4.7^o^C/min and -1.5^o^C/min) and with various cryoprotectant agents (n=8). Abbreviations: Me2SO = dimethyl sulfoxide, PG = propylene glycol, EG = ethylene glycol. Note letters identify significant subsets (*p* < 0.05 Wilcoxin pairwise comparison).

Table S2-A: Comparison of *Tripneustes gratilla* survivorship distributions between cryoprotectant agents and cooling rate treatments. Abbreviations: Me2SO = dimethyl sulfoxide, PG = propylene glycol, EG = ethylene glycol, FSW = filtered seawater.

| **Between** | **df** | **Wilcoxin** | | **Survivorship Verdict** |
| --- | --- | --- | --- | --- |
|  |  | **Statistic** | **P** |  |
| All treatments | 12 | 93.3653 | <0.0001 |  |
| 1M Me2SO, -15.2C/min vs 1M Me2SO, -1.5C/min | 1 | 3.53361 | 0.0004 | 1M Me2SO, -15.2C/min > 1M Me2SO, -1.5C/min |
| 1M Me2SO, -9.7C/min vs 1M Me2SO, -1.5C/min | 1 | 3.53361 | 0.0004 | 1M Me2SO, -9.7C/min > 1M Me2SO, -1.5C/min |
| 1M Me2SO, -9.7C/min vs 1M Me2SO, -4.7C/min | 1 | 3.53361 | 0.0004 | 1M Me2SO, -9.7C/min > 1M Me2SO, -4.7C/min |
| 1M EG, -15.2C/min vs 1MMe2SO, -1.5C/min | 1 | 3.53361 | 0.0004 | 1M EG, -15.2C/min > 1MMe2SO, -1.5C/min |
| 1M EG, -15.2C/min vs 1MMe2SO, -4.7C/min | 1 | 3.53361 | 0.0004 | 1M EG, -15.2C/min > 1MMe2SO, -4.7C/min |
| 1M EG, -15.2C/min vs 1M EG, -1.5C/min | 1 | 3.53361 | 0.0004 | 1M EG, -15.2C/min > 1M EG, -1.5C/min |
| 1M EG, -9.7C/min vs 1MMe2SO, -1.5C/min | 1 | 3.53361 | 0.0004 | 1M EG, -9.7C/min > 1MMe2SO, -1.5C/min |
| 1M EG, -9.7C/min vs 1MMe2SO, -4.7C/min | 1 | 3.53361 | 0.0004 | 1M EG, -9.7C/min > 1MMe2SO, -4.7C/min |
| 1M EG, -9.7C/min vs 1M EG, -1.5C/min | 1 | 3.53361 | 0.0004 | 1M EG, -9.7C/min > 1M EG, -1.5C/min |
| 1M EG, -9.7C/min vs 1M EG, -4.7C/min | 1 | 3.53361 | 0.0004 | 1M EG, -9.7C/min > 1M EG, -4.7C/min |
| 1M Me2SO, -9.7C/min vs 1M Me2SO, -15.2C/min | 1 | 3.09812 | 0.0019 | 1M Me2SO, -9.7C/min > 1M Me2SO, -15.2C/min |
| 1M PG, -4.7C/min vs 1M Me2SO, -1.5C/min | 1 | 3.18251 | 0.0015 | 1M PG, -4.7C/min > 1M Me2SO, -1.5C/min |
| 1M PG, -4.7C/min vs 1M Me2SO, -4.7C/min | 1 | 3.18251 | 0.0015 | 1M PG, -4.7C/min > 1M Me2SO, -4.7C/min |
| 1M PG, -4.7C/min vs 1M EG, -1.5C/min | 1 | 3.18251 | 0.0015 | 1M PG, -4.7C/min > 1M EG, -1.5C/min |
| 1M PG, -4.7C/min vs 1M EG, -4.7C/min | 1 | 3.18251 | 0.0015 | 1M PG, -4.7C/min > 1M EG, -4.7C/min |
| 1M PG, -4.7C/min vs 1M PG, -1.5C/min | 1 | 3.18251 | 0.0015 | 1M PG, -4.7C/min > 1M PG, -1.5C/min |
| 1M PG, -4.7C/min vs 1M PG, -15.2C/min | 1 | 3.18251 | 0.0015 | 1M PG, -4.7C/min > 1M PG, -15.2C/min |
| FSW vs 1M Me2SO, -1.5C/min | 1 | 3.35978 | 0.0008 | FSW > 1M Me2SO, -1.5C/min |
| FSW vs 1M Me2SO, -4.7C/min | 1 | 3.35978 | 0.0008 | FSW > 1M Me2SO, -4.7C/min |
| FSW vs 1M EG, -1.5C/min | 1 | 3.35978 | 0.0008 | FSW > 1M EG, -1.5C/min |
| FSW vs 1M EG, -15.2C/min | 1 | 3.03384 | 0.0024 | FSW > 1M EG, -15.2C/min |
| FSW vs 1M EG, -4.7C/min | 1 | 3.35978 | 0.0008 | FSW > 1M EG, -4.7C/min |
| FSW vs 1M EG, -9.7C/min | 1 | 3.03384 | 0.0024 | FSW >1M EG, -9.7C/min |
| FSW vs 1M Me2SO, 15.2C/min | 1 | 2.51744 | 0.0118 | FSW > 1M Me2SO, 15.27C/min |
| 1M PG, -4.7C/min vs 1M EG, -15.2 C/min | 1 | 1.52280 | 0.1270 | NOT SIGNFICANT |
| 1M EG, -9.7C/min vs 1M EG, -15.2C/min | 1 | 0.68264 | 0.2271 | NOT SIGNIFICANT |
| 1M PG, -4.7C/min vs 1M EG, -9.7C/min | 1 | 0.68476 | 0.4948 | NOT SIGNIFICANT |
| 1M PG, -9.7C/min vs 1M Me2SO, -1.5C/min | 1 | 0.87500 | 0.3816 | NOT SIGNFICANT |
| 1M PG, -9.7C/min vs 1M Me2SO, -4.7C/min | 1 | 0.87500 | 0.3816 | NOT SIGNFICANT |
| 1M PG, -9.7C/min vs 1M EG, -1.5C/min | 1 | 0.87500 | 0.3816 | NOT SIGNFICANT |
| 1M PG, -9.7C/min vs 1M EG, -4.7C/min | 1 | 0.87500 | 0.3816 | NOT SIGNFICANT |
| 1M PG, -9.7C/min vs 1M PG, -1.5C/min | 1 | 0.87500 | 0.3816 | NOT SIGNFICANT |
| 1M PG, -9.7C/min vs 1M PG, -15.2C/min | 1 | 0.87500 | 0.3816 | NOT SIGNFICANT |
| FSW vs 1M Me2SO, -9.7C/min | 1 | 0.32275 | 0.7469 | NOT SIGNIFICANT |
| 1M Me2SO, -4.7C/min vs 1M Me2SO, -1.5C/min | 1 | 0 | 1 | NOT SIGNIFICANT |
| 1M EG, -1.5C/min vs 1M Me2SO, -1.5C/min | 1 | 0 | 1 | NOT SIGNIFICANT |
| 1M EG, -1.5C/min vs 1M Me2SO, -4.7C/min | 1 | 0 | 1 | NOT SIGNIFICANT |
| 1M EG, -4.7C/min vs 1M Me2SO, -1.5C/min | 1 | 0 | 1 | NOT SIGNIFICANT |
| 1M EG, -4.7C/min vs 1M Me2SO, -4.7C/min | 1 | 0 | 1 | NOT SIGNIFICANT |
| 1M EG, -4.7C/min vs 1M EG, -1.5C/min | 1 | 0 | 1 | NOT SIGNIFICANT |
| 1M PG, -1.5C/min vs 1M Me2SO, -1.5C/min | 1 | 0 | 1 | NOT SIGNIFICANT |
| 1M PG, -1.5C/min vs 1M Me2SO, -4.7C/min | 1 | 0 | 1 | NOT SIGNIFICANT |
| 1M PG, -1.5C/min vs 1M EG, -1.5C/min | 1 | 0 | 1 | NOT SIGNIFICANT |
| 1M PG, -1.5C/min vs 1M EG, -4.7C/min | 1 | 0 | 1 | NOT SIGNIFICANT |
| 1M PG, -15.2C/min vs 1M Me2SO, -1.5C/min | 1 | 0 | 1 | NOT SIGNIFICANT |
| 1M PG, -15.2C/min vs 1M Me2SO, -4.7C/min | 1 | 0 | 1 | NOT SIGNIFICANT |
| 1M PG, -15.2C/min vs 1M EG, -1.5C/min | 1 | 0 | 1 | NOT SIGNIFICANT |
| 1M PG, -15.2C/min vs 1M EG, -4.7C/min | 1 | 0 | 1 | NOT SIGNIFICANT |
| 1M PG, -15.2C/min vs 1M PG, -1.5C/min | 1 | 0 | 1 | NOT SIGNIFICANT |
| 1M PG, -4.7C/min vs 1M Me2SO, -15.2C/min | 1 | -0.36757 | 0.7132 | NOT SIGNIFICANT |
| 1M EG, -9.7C/min vs 1M Me2SO, -15.2C/min | 1 | -1.36628 | 0.1719 | NOT SIGNIFICANT |
| 1M PG, -4.7C/min vs FSW | 1 | -2.25924 | 0.0239 | 1M PG, -4.7C/min < FSW |
| 1M EG, -15.2C/min vs 1M Me2SO, -15.2C/min | 1 | -2.15293 | 0.0313 | 1M EG, -15.2C/min < 1M Me2SO, -15.2C/min |
| 1M PG, -4.7C/min vs 1M Me2SO, -9.7C/min | 1 | -2.780476 | 0.0054 | 1M PG, -4.7C/min < 1M Me2SO, -9.7C/min |
| 1M PG, -9.7C/min vs 1M PG, -4.7C/min | 1 | -3.02880 | 0.0025 | 1M PG, -9.7C/min < 1M PG, -4.7C/min |
| 1M PG, -1.5C/min vs FSW | 1 | -3.35978 | 0.0008 | 1M PG, -1.5C/min < FSW |
| 1M PG, -15.2C/min vs FSW | 1 | -3.35978 | 0.0008 | 1M PG, -15.2C/min < FSW |
| 1M PG, -9.7C/min vs FSW | 1 | -3.23975 | 0.0012 | 1M PG, -9.7C/min < FSW |
| 1M Me2SO, -4.7C/min vs 1M Me2SO, -15.2C/min | 1 | -3.53361 | 0.0004 | 1M Me2SO, -4.7C/min < 1M Me2SO, -15.2C/min |
| 1M EG, -1.5C/min vs 1M Me2SO, -15.2C/min | 1 | -3.53361 | 0.0004 | 1M EG, -1.5C/min < 1M Me2SO, -15.2C/min |
| 1M EG, -1.5C/min vs 1M Me2SO, -9.7C/min | 1 | -3.53361 | 0.0004 | 1M EG, -1.5C/min < 1M Me2SO, -9.7C/min |
| 1M EG, -15.2C/min vs 1M Me2SO, -9.7C/min | 1 | -3.30816 | 0.0009 | 1M EG, -15.2C/min < 1M Me2SO, -9.7C/min |
| 1M EG, -4.7C/min vs 1M Me2SO, -15.2C/min | 1 | -3.53361 | 0.0004 | 1M EG, -4.7C/min < 1M Me2SO, -15.2C/min |
| 1M EG, -4.7C/min vs 1M Me2SO, -9.7C/min | 1 | -3.53361 | 0.0004 | 1M EG, -4.7C/min < 1M Me2SO, -9.7C/min |
| 1M EG, -4.7C/min vs 1M EG, -15.2C/min | 1 | -3.53361 | 0.0004 | 1M EG, -4.7C/min < 1M EG, -15.2C/min |
| 1M EG, -9.7C/min vs 1M Me2SO, -9.7C/min | 1 | -3.30816 | 0.0009 | 1M EG, -9.7C/min < 1M Me2SO, -9.7C/min |
| 1M PG, -1.5C/min vs 1M Me2SO, -15.2C/min | 1 | -3.53361 | 0.0004 | 1M PG, -1.5C/min < 1M Me2SO, -15.2C/min |
| 1M PG, -1.5C/min vs 1M Me2SO, -9.7C/min | 1 | -3.53361 | 0.0004 | 1M PG, -1.5C/min < 1M Me2SO, -9.7C/min |
| 1M PG, -1.5C/min vs 1M EG, -15.2C/min | 1 | -3.53361 | 0.0004 | 1M PG, -1.5C/min < 1M EG, -15.2C/min |
| 1M PG, -1.5C/min vs 1M EG, -9.7C/min | 1 | -3.53361 | 0.0004 | 1M PG, -1.5C/min < 1M EG, -9.7C/min |
| 1M PG, -15.2C/min vs 1M Me2SO, -1.52C/min | 1 | -3.53361 | 0.0004 | 1M PG, -15.2C/min < 1M Me2SO, -1.52C/min |
| 1M PG, -15.2C/min vs 1M Me2SO, -9.7C/min | 1 | -3.53361 | 0.0004 | 1M PG, -15.2C/min < 1M Me2SO, -9.7C/min |
| 1M PG, -15.2C/min vs 1M EG, -15.2C/min | 1 | -3.53361 | 0.0004 | 1M PG, -15.2C/min < 1M EG, -15.2C/min |
| 1M PG, -15.2C/min vs 1M EG, -9.7C/min | 1 | -3.53361 | 0.0004 | 1M PG, -15.2C/min < 1M EG, -9.7C/min |
| 1M PG, -9.7C/min vs 1M Me2SO, -15.2C/min | 1 | -3.45342 | 0.0006 | 1M PG, -9.7C/min < 1M Me2SO, -15.2C/min |
| 1M PG, -9.7C/min vs 1M Me2SO, -9.7C/min | 1 | -3.45342 | 0.0006 | 1M PG, -9.7C/min < 1M Me2SO, -9.7C/min |
| 1M PG, -9.7C/min vs 1M EG, -15.2C/min | 1 | -3.45342 | 0.0006 | 1M PG, -9.7C/min < 1M EG, -15.2C/min |
| 1M PG, -4.7C/min vs 1M EG, -9.7C/min | 1 | -3.45342 | 0.0006 | 1M PG, -4.7C/min < 1M EG, -9.7C/min |

Table S2-B: Comparison of *Tripneustes gratilla* development distributions to the 4-arm pluteus stage between cryoprotectant agents and cooling rate treatments. Abbreviations: Me2SO = dimethyl sulfoxide, PG = propylene glycol, EG = ethylene glycol, FSW = filtered seawater.

| **Between** | **df** | **Wilcoxin** | | **Survivorship Verdict** |
| --- | --- | --- | --- | --- |
|  |  | **Statistic** | **P** |  |
| All treatments | 12 | 77.6065 | <0.0001 |  |
| 1M Me2SO, -9.7C/min vs 1M Me2SO, -1.5C/min | 1 | 3.53361 | 0.0004 | 1M Me2SO, -9.7C/min > 1M Me2SO, -1.5C/min |
| 1M Me2SO, -9.7C/min vs 1M Me2SO, -4.7C/min | 1 | 3.53361 | 0.0004 | 1M Me2SO, -9.7C/min > 1M Me2SO, -4.7C/min |
| FSW vs 1M Me2SO, -1.5C/min | 1 | 3.25978 | 0.0008 | FSW > 1M Me2SO, -1.5C/min |
| FSW vs 1M Me2SO, -4.7C/min | 1 | 3.25978 | 0.0008 | FSW > 1MMe2SO, -4.7C/min |
| FSW vs 1M EG, -1.5C/min | 1 | 3.25978 | 0.0008 | FSW > 1M EG, -1.5C/min |
| FSW vs 1M EG, -15.2C/min | 1 | 3.25978 | 0.0008 | FSW > 1M EG, -15.2C/min |
| FSW vs 1M EG, -4.7C/min | 1 | 3.25978 | 0.0008 | FSW > 1M EG, -4.7C/min |
| FSW vs 1M EG, -9.7C/min | 1 | 2.80665 | 0.0050 | FSW > 1M EG, -9.7C/min |
| 1M Me2SO, -15.2C/min vs 1M Me2SO, -1.5C/min | 1 | 2.83593 | 0.0046 | 1M Me2SO, -15.2C/min > 1M Me2SO, -1.5C/min |
| FSW vs 1M Me2SO, -15.2C/min | 1 | 2.39097 | 0.0168 | FSW > 1M Me2SO, -15.2C/min |
| 1M PG, -4.7C/min vs 1M Me2SO, -1.5C/min | 1 | 2.48993 | 0.0128 | 1M PG, -4.7C/min > 1M Me2SO, -1.5C/min |
| 1M PG, -4.7C/min vs 1M Me2SO, -4.7C/min | 1 | 2.48993 | 0.0128 | 1M PG, -4.7C/min > 1M Me2SO, -4.7C/min |
| 1M PG, -4.7C/min vs 1M EG, -1.5C/min | 1 | 2.48993 | 0.0128 | 1M PG, -4.7C/min > 1M EG, -1.5C/min |
| 1M PG, -4.7C/min vs 1M EG, -15.2C/min | 1 | 2.48993 | 0.0128 | 1M PG, -4.7C/min > 1M EG, -15.2C/min |
| 1M PG, -4.7C/min vs 1M EG, -4.7C/min | 1 | 2.48993 | 0.0128 | 1M PG, -4.7C/min > 1M EG, -4.7C/min |
| 1M PG, -4.7C/min vs 1M PG, -1.5C/min | 1 | 2.48993 | 0.0128 | 1M PG, -4.7C/min > 1M PG, -1.5C/min |
| 1M PG, -4.7C/min vs 1M PG, -15.2C/min | 1 | 2.48993 | 0.0128 | 1M PG, -4.7C/min > 1M PG, -15.2C/min |
| FSW vs 1M Me2SO, -9.7C/min | 1 | 1.87194 | 0.0612 | FSW > 1M Me2SO, -9.7C/min |
| 1M EG, -9.7C/min vs 1M Me2SO, -1.5C/min | 1 | 2.13852 | 0.0325 | 1M EG, -9.7C/min > 1M Me2SO, -1.5C/min |
| 1M EG, -9.7C/min vs 1M Me2SO, -4.7C/min | 1 | 2.13852 | 0.0325 | 1M EG, -9.7C/min > 1M Me2SO, -4.7C/min |
| 1M EG, -9.7C/min vs 1M EG, -1.5C/min | 1 | 2.13852 | 0.0325 | 1M EG, -9.7C/min > 1M EG, -1.5C/min |
| 1M EG, -9.7C/min vs 1M EG, -15.2C/min | 1 | 2.13852 | 0.0325 | 1M EG, -9.7C/min > 1M EG, -15.2C/min |
| 1M EG, -9.7C/min vs 1M EG, -4.7C/min | 1 | 2.13852 | 0.0325 | 1M EG, -9.7C/min > 1M EG, -4.7C/min |
| 1M Me2SO, -9.7C/min vs 1M Me2SO, -15.2C/min | 1 | 1.57764 | 0.1146 | NOT SIGNFICANT |
| 1M PG, -4.7C/min vs 1M EG, -9.7 C/min | 1 | 0.93187 | 0.3514 | NOT SIGNFICANT |
| 1M Me2SO, -4.7C/min vs 1M Me2SO, -1.5C/min | 1 | 0 | 1 | NOT SIGNIFICANT |
| 1M EG, -1.5C/min vs 1M Me2SO, -1.5C/min | 1 | 0 | 1 | NOT SIGNIFICANT |
| 1M EG, -1.5C/min vs 1M Me2SO, -4.7C/min | 1 | 0 | 1 | NOT SIGNFICANT |
| 1M EG, -15.2C/min vs 1M Me2SO, -1.5C/min | 1 | 0 | 1 | NOT SIGNFICANT |
| 1M EG, -15.2C/min vs 1M Me2SO, -4.7C/min | 1 | 0 | 1 | NOT SIGNFICANT |
| 1M EG, -15.2C/min vs 1M EG, -1.5C/min | 1 | 0. | 1 | NOT SIGNFICANT |
| 1M EG, -4.7C/min vs 1M Me2SO, -1.5C/min | 1 | 0. | 1 | NOT SIGNFICANT |
| 1M EG, -4.7C/min vs 1M Me2SO, -4.7C/min | 1 | 0. | 1 | NOT SIGNFICANT |
| 1M EG, -4.7C/min vs 1M EG, -1.5C/min | 1 | 0. | 1 | NOT SIGNFICANT |
| 1M EG, -4.7C/min vs 1M EG, -15.2C/min | 1 | 0 | 1 | NOT SIGNIFICANT |
| 1M PG, -1.5C/min vs 1M Me2SO, -1.5C/min | 1 | 0 | 1 | NOT SIGNIFICANT |
| 1M PG, -1.5C/min vs 1M Me2SO, -4.7C/min | 1 | 0 | 1 | NOT SIGNIFICANT |
| 1M PG, -1.5C/min vs 1M EG, -1.5C/min | 1 | 0 | 1 | NOT SIGNIFICANT |
| 1M PG, -1.5C/min vs 1M EG, -15.2C/min | 1 | 0 | 1 | NOT SIGNIFICANT |
| 1M PG, -1.5C/min vs 1M EG, -4.7C/min | 1 | 0 | 1 | NOT SIGNIFICANT |
| 1M PG, -15.2C/min vs 1M Me2SO, -1.5C/min | 1 | 0 | 1 | NOT SIGNIFICANT |
| 1M PG, -15.2C/min vs 1M Me2SO, -4.7C/min | 1 | 0 | 1 | NOT SIGNIFICANT |
| 1M PG, -15.2C/min vs 1M EG, -1.5C/min | 1 | 0 | 1 | NOT SIGNIFICANT |
| 1M PG, -15.2C/min vs 1M EG, -15.2C/min | 1 | 0 | 1 | NOT SIGNIFICANT |
| 1M PG, -15.2C/min vs 1M EG, -4.7C/min | 1 | 0 | 1 | NOT SIGNIFICANT |
| 1M PG, -15.2C/min vs 1M PG, -1.5C/min | 1 | 0 | 1 | NOT SIGNIFICANT |
| 1M PG, -9.7C/min vs 1M Me2SO, -1.5C/min | 1 | 0 | 1 | NOT SIGNIFICANT |
| 1M PG, -9.7C/min vs 1M Me2SO, -4.7C/min | 1 | 0 | 1 | NOT SIGNIFICANT |
| 1M PG, -9.7C/min vs 1M EG, -1.5C/min | 1 | 0 | 1 | NOT SIGNIFICANT |
| 1M PG, -9.7C/min vs 1M EG, -15.2C/min | 1 | 0 | 1 | NOT SIGNIFICANT |
| 1M PG, -9.7C/min vs 1M EG, -4.7C/min | 1 | 0 | 1 | NOT SIGNIFICANT |
| 1M PG, -9.7C/min vs 1M PG, -1.5C/min | 1 | 0 | 1 | NOT SIGNIFICANT |
| 1M PG, -9.7C/min vs 1M PG, -15.2C/min | 1 | 0 | 1 | NOT SIGNIFICANT |
| 1M PG, -4.7C/min vs 1M Me2SO, -15.2C/min | 1 | -0.47970 | 0.06314 | NOT SIGNIFICANT |
| 1M EG, -9.7C/min vs 1M Me2SO, -15.2C/min | 1 | -1.56358 | 0.1179 | NOT SIGNIFICANT |
| 1M PG, -1.5C/min vs 1M EG, -9.7C/min | 1 | -2.13852 | 0.0325 | 1M PG, -1.5C/min < 1M EG, -9.7C/min |
| 1M PG, -15.2C/min vs 1M EG, -9.7C/min | 1 | -2.13852 | 0.0325 | 1M PG, -15.2C/min < 1M EG, -9.7C/min |
| 1M PG, -9.7C/min vs 1M EG, -9.7C/min | 1 | -2.13852 | 0.0325 | 1M PG, -9.7C/min < 1M EG, -9.7C/min |
| 1M PG, -9.7C/min vs 1M PG, -4.7C/min | 1 | -2.48993 | 0.0128 | 1M PG, -9.7C/min < 1M PG, -4.7C/min |
| 1M PG, -4.7C/min vs 1M Me2SO, -9.7C/min | 1 | -2.15929 | 0.0308 | 1M PG, -4.7C/min < 1M Me2SO, -9.7C/min |
| 1M PG, -4.7C/min vs FSW | 1 | -2.52858 | 0.0115 | 1M PG, -4.7C/min < FSW |
| 1M Me2SO, -4.7C/min vs 1M Me2SO, -15.2C/min | 1 | -2.83593 | 0.0046 | 1M Me2SO, -4.7C/min < 1M Me2SO, -15.2C/min |
| 1M EG, -1.5C/min vs 1M Me2SO, -15.2C/min | 1 | -2.83593 | 0.0046 | 1M EG, -1.5C/min < 1M Me2SO, -15.2C/min |
| 1M EG, -15.2C/min vs 1M Me2SO, -15.2C/min | 1 | -2.83593 | 0.0046 | 1M EG, -15.2C/min < 1M Me2SO, -15.2C/min |
| 1M EG, -4.7C/min vs 1M Me2SO, -15.2C/min | 1 | -2.83593 | 0.0046 | 1M EG, -4.7C/min < 1M Me2SO, -15.2C/min |
| 1M EG, -9.7C/min vs 1M Me2SO, -9.7C/min | 1 | -2.48634 | 0.0129 | 1M EG, -9.7C/min < 1M Me2SO, -9.7C/min |
| 1M PG, -1.5C/min vs 1M Me2SO, -15.2C/min | 1 | -2.83593 | 0.0046 | 1M PG, -1.5C/min < 1M Me2SO, -15.2C/min |
| 1M PG, -15.2C/min vs 1M Me2SO, -15.2C/min | 1 | -2.83593 | 0.0046 | 1M PG, -15.2C/min < 1M Me2SO, -15.2C/min |
| 1M PG, -9.7C/min vs 1M Me2SO, -15.2C/min | 1 | -2.83593 | 0.0046 | 1M PG, -9.7C/min < 1M Me2SO, -15.2C/min |
| 1M PG, -1.5C/min vs FSW | 1 | -3.35978 | 0.0008 | 1M PG, -1.5C/min < FSW |
| 1M PG, -15.2C/min vs FSW | 1 | -3.35978 | 0.0008 | 1M PG, -15.2C/min < FSW |
| 1M PG, -9.7C/min vs FSW | 1 | -3.35978 | 0.0008 | 1M PG, -9.7C/min < FSW |
| 1M EG, -1.5C/min vs 1M Me2SO, -9.7C/min | 1 | -3.53361 | 0.0004 | 1M EG, -1.5C/min < 1M Me2SO, -9.7C/min |
| 1M EG, -15.2C/min vs 1M Me2SO, -9.7C/min | 1 | -3.53361 | 0.0004 | 1M EG, -15.2C/min < 1M Me2SO, -9.7C/min |
| 1M EG, -4.7C/min vs 1M Me2SO, -9.7C/min | 1 | -3.53361 | 0.0004 | 1M EG, -4.7C/min < 1M Me2SO, -9.7C/min |
| 1M PG, -1.5C/min vs 1M Me2SO, -9.7C/min | 1 | -3.53361 | 0.0004 | 1M PG, -1.5C/min < 1M Me2SO, -9.7C/min |
| 1M PG, -15.2C/min vs 1M Me2SO, -9.7C/min | 1 | -3.53361 | 0.0004 | 1M PG, -15.2C/min < 1M Me2SO, -9.7C/min |
| 1M PG, -9.7C/min vs 1M Me2SO, -9.7C/min | 1 | -3.53361 | 0.0004 | 1M PG, -9.7C/min < 1M Me2SO, -9.7C/min |

2-B) Testing the addition of trehalose to dimethyl sulfoxide

Figure S5: Kaplan-Meier product-limit survivorship curves comparing *Tripneustes gratilla* embryos cryopreserved with different 1M Me2SO, 1M Me2SO + 0.05M TRE and 1M Me2SO + 0.1M TRE (n=800). Abbreviations: Me2SO = dimethyl sulfoxide, TRE = trehalose.

Table S3: Comparison of *Tripneustes gratilla* survivorship distributions between 1M Me2SO, 1M Me2SO + 0.05M TRE and 1M Me2SO + 0.1M TRE. Abbreviations: Me2SO = dimethyl sulfoxide, TRE = trehalose.

| **Between** | **df** | **Wilcoxin** | | **Survivorship Verdict** |
| --- | --- | --- | --- | --- |
|  |  | **Statistic** | **P** |  |
| All Treatments | 3 | 15.6338 | 0.0013 |  |
| FSW vs 1M Me2SO + 0.1M TRE | 1 | 2.90955 | 0.0036 | FSW > 1M Me2SO + 0.1M TRE |
| FSW vs 1M Me2SO + 0.05M TRE | 1 | 2.16894 | 0.0301 | FSW > 1M Me2SO + 0.05M TRE |
| FSW vs 1M Me2SO | 1 | 1.85153 | 0.0641 | NOT SIGNFICANT |
| 1M Me2SO + 0.1M TRE vs 1M Me2SO + 0.05M TRE | 1 | -1.31276 | 0.1893 | NOT SIGNFICANT |
| 1M Me2SO + 0.05M TRE vs 1M Me2SO | 1 | -1.52280 | 0.1278 | NOT SIGNFICANT |
| 1M Me2SO + 0.1M TRE vs 1M Me2SO | 1 | -3.20314 | 0.0014 | 1M Me2SO + 0.1M TRE < 1M Me2SO |

Figure S6: Proportion of *Tripneustes gratilla* offspring at various stages of development (gastrula, prism, 2-arm pluteus, 4-arm pluteus),72 hours post thawing from cryopreservation with dimethyl sulfoxide mixed with trehalose (n=8). Abbreviations: TRE = trehalose, Me2SO = dimethyl sulfoxide. Note letters identify significant subsets (*p* < 0.05 Wilcoxin pairwise comparison).

Trehalose supplementation of cryoprotocols was examined at two concentrations. The differences in survival rates between cryopreserved protocols were found to be significant (*p* < 0.05, Kaplan-Meier survivorship curves, Figure S5). The addition of 0.1M TRE and 0.05M TRE to 1M Me2SO significantly decreased survivorship to 33.9% and 58.6%, respectively, compared with a survivorship of 88.5% for embryos from the control treatments (Figure S5, S6) (*p* < 0.05, Wilcoxin pairwise comparison) (Table S3). No significant difference in survivorship between embryos cryopreserved with 1M Me2SO and our control embryos, reared in FSW without being cryopreserved, were found (survivorships were 85.5±3.2% and 88.5±8.6%, respectively, Figure S2-a,b,c) (*p* > 0.05, Wilcoxin pairwise comparison) (Table S3). Nearly all surviving embryos cryopreserved with TRE remained stunted in the gastrula stage, whereas only 15.2 ± 4.4% of embryos treated with 1M Me2SO were stunted in the gastrula stage (Figure S2-c). An average of 35.5 ± 8.2% of embryos cryopreserved with 1M Me2SO continued to develop to the 4-arm pluteus stage within 72 hours of thawing (Figure S2-c).

*Supplemental Discussion for this experiment:*

Non-permeant cryoprotectants are sometimes used because they serve as osmotic buffers, thereby reducing the osmotic shock and toxicity from exposure to permeant cryoprotectants [18,77,54,1,8,52]. Additionally, non-permeant cryoprotectant saccharides (sucrose, trehalose, glucose) form hydrogen bonds with the phospholipid polar groups in membrane bilayers which can help stabilize the membrane during freezing and thawing [70]. Even so, the addition of non-permeant cryoprotectant does not always improve long-term survival of cryopreserved sea urchin larvae [2]. To determine if the addition of the non-permeant cryoprotection agent, trehalose, would provide any benefits to the freezing of *T. gratilla* embryos, the most effective protocol from the previous experiment (1M Me2SO cooled at -9.7**°**C/min) was supplemented with TRE. The addition of TRE did not improve survivorship or developmental rates in *T. gratilla* embryo cryopreservation, a conclusion that seems consistent with our toxicity trials (Figure S1 & S2). When compared to the control treatments, the addition of TRE to the cryopreservation protocol simultaneously decreased survivorship, in proportion with concentration of trehalose added to the protocol (1M Me2SO + 0.1M TRE & 1M Me2SO + 0.05M TRE) and stunted development in the embryonic stage (Figure S5 & S6) (*p* < 0.05 Wilcoxin pairwise comparison) (Table S3) (results in supplemental information). However, treatments that cryopreserved embryos using only 1M Me2SO had survivorships that were not significantly different from the control treatments (Figure S5 & S6) (*p* > 0.05 Wilcoxin pairwise comparison) (Table S3) (results in supplemental information). Although some studies have reported an increase in the efficacy of their cryopreservation protocols with the addition of trehalose [13,49,53,8], others have reported that this non-permeant cryoprotectant agent did not improve long-term survival of their cryopreserved specimens [2].

A noteworthy result from this experiment was that the samples cryopreserved in 1M Me2SO, did not have a significantly lower survival rate than our controlled samples, which were simply incubated in FSW (Figure S5 & S6) (*p* > 0.05 Wilcoxin pairwise comparison) (Table S3)(results in supplemental information). Albeit, the control treatments tended to have a higher proportion of larvae that advanced to the 4-arm pluteus stage than the cryopreserved treatments, for these experiments.

3) Cryopreservation at various larval stages

Figure S7: Kaplan-Meier product-limit survivorship curves comparing the cryopreservation of *Tripneustes gratilla* from various early developmental stages (n=800).

Figure S8: Developmental stages of *Tripneustes gratilla* offspring cryopreserved (with 1M Me2SO at -6.5**°**C/min) at various stages of development (fertilized egg, blastula, gastrula and pluteus), 72 hours post thawing (n=8). Note letters identify significant subsets (*p* < 0.05 Wilcoxin pairwise comparison).

Table S4: Comparison of survivorship distributions between cryopreserved *Tripneustes gratilla* fertilized eggs, blastula, gastrula and 4-arm pluteus larvae

| **Between** | **df** | **Wilcoxin** | | **Survivorship Verdict** |
| --- | --- | --- | --- | --- |
|  |  | **Statistic** | **P** |  |
| All Treatments | 4 | 33.3150 | <0.0001 |  |
| Control vs 4-arm Plut | 1 | 3.45342 | 0.0006 | Control > 4-arm Plut |
| Gastrula vs 4-arm Plut | 1 | 3.30816 | 0.0009 | Gastrula > 4-arm Plut |
| Gastrula vs Fert Egg | 1 | 3.53361 | 0.0004 | Gastrula > Fert Egg |
| Control vs Blastula | 1 | 3.01489 | 0.0026 | Control > Blastula |
| Gastrula vs Blastula | 1 | 1.94289 | 0.0520 | NOT SIGNFICANT |
| Blastula vs 4-arm Put | 1 | 1.83787 | 0.0661 | NOT SIGNIFICANT |
| Gastrula vs Control | 1 | -2.68599 | 0.0072 | Gastrula < Control |
| Fertilized Egg vs 4-arm Plut | 1 | -3.53361 | 0.0004 | Fertilized Egg < 4-arm Plut |
| Fertilized Egg vs Blastula | 1 | -3.53361 | 0.0004 | Fertilized Egg < Blastula |
| Fert Egg vs Control | 1 | -3.71231 | 0.0002 | Fert Egg < Control |

Table S5: Comparison of developmental distributions to the 4-arm pluteus stage between cryopreserved *Tripneustes gratilla* fertilized eggs, blastula, gastrula and 4-arm pluteus larvae

| **Between** | **df** | **Wilcoxin** | | **Survivorship Verdict** |
| --- | --- | --- | --- | --- |
|  |  | **Statistic** | **P** |  |
| All Treatments | 4 | 30.8942 | <0.0001 |  |
| Control vs 4-arm Plut | 1 | 3.45342 | 0.0006 | Control > 4-arm Plut |
| Gastrula vs 4-arm Plut | 1 | 2.04791 | 0.0406 | Gastrula > 4-arm Plut |
| Gastrula vs Fert Egg | 1 | 3.53361 | 0.0004 | Gastrula > Fert Egg |
| Control vs Blastula | 1 | 3.23415 | 0.0012 | Control > Blastula |
| Gastrula vs Blastula | 1 | 1.52280 | 0.1278 | NOT SIGNFICANT |
| Blastula vs 4-arm Put | 1 | -0.36757 | 0.7132 | NOT SIGNIFICANT |
| Gastrula vs Control | 1 | -3.01489 | 0.0026 | Gastrula < Control |
| Fertilized Egg vs 4-arm Plut | 1 | -3.53361 | 0.0004 | Fertilized Egg < 4-arm Plut |
| Fertilized Egg vs Blastula | 1 | -3.53361 | 0.0004 | Fertilized Egg < Blastula |
| Fert Egg vs Control | 1 | -3.71231 | 0.0002 | Fert Egg < Control |
